# How important is CRISPR-Cas for protecting natural populations of bacteria against infections by mobile genetic elements?

**DOI:** 10.1101/2020.02.05.935965

**Authors:** Edze Westra, Bruce Levin

## Abstract

Articles on CRISPR commonly open with some variant of the phrase ‘these short-palindromic repeats and their associated endonucleases (Cas) are an adaptive immune system that exists to protect bacteria and archaea from viruses and infections with other mobile genetic elements’. There is an abundance of genomic data consistent with the hypothesis that CRISPR plays this role in natural populations of bacteria and archaea, and experimental demonstrations with a few species of bacteria and their phage and plasmids show that CRISPR-Cas systems can play this role *in vitro*. Not at all clear are the ubiquity, magnitude and nature of the contribution of CRISPR-Cas systems to the ecology and evolution of natural populations of microbes, and the strength of selection mediated by different types of phage and plasmids to the evolution and maintenance of CRISPR-Cas systems. In this perspective, with the aid of heuristic mathematical-computer simulation models, we explore the *a priori* conditions under which exposure to lytic and temperate phage and conjugative plasmids will select for and maintain CRISPR-Cas systems in populations of bacteria and archaea. We review the existing literature addressing these ecological and evolutionary questions and highlight the experimental and other evidence needed to fully understand the conditions responsible for the evolution and maintenance of CRISPR-Cas systems and the contribution of these systems to the ecology and evolution of bacteria, archaea and the mobile genetic elements that infect them.

**Significance:** There is no question about the importance and utility of CRISPR-Cas for editing and modifying genomes. On the other hand, the mechanisms responsible for the evolution and maintenance of these systems and the magnitude of their importance to the ecology and evolution of bacteria, archaea and their infectious DNAs, are not at all clear. With the aid of heuristic mathematical – computer simulation models and reviews of the existing literature, we raise questions that have to be answered to elucidate the contribution of selection – mediated by phage and plasmids – to the evolution and maintenance of this adaptive immune system and its consequences for the ecology and evolution of prokaryotes and their viruses and plasmids.

## Introduction

In 1987, a study by Ishino and colleagues aimed at analyzing the nucleotide sequence of the *iap* gene (isozyme-converting alkaline phosphatase) in *Escherichia coli* serendipitously led to the first-ever description of a CRISPR array. Specifically, they identified 14 repetitive sequences of 29 base pairs each at the 3’ end of the *iap* gene, that were interspersed by 32–33 bp variable sequences (1, 2). Over the next years, identification of CRISPR arrays in other Gram-negative bacteria, Gram-positives and in Archaea (3-7) triggered a quest to identify their biological function (reviewed in (8)). Meanwhile, as more whole-genome sequences became available and CRISPR detection algorithms were developed, it became clear that these arrays of repeating sequences are common in prokaryotes, with estimated frequencies of approximately 30-40% in Bacteria and 90% in the Archaea (9-13), with clear variation between phyla (14). An important step in understanding the function of CRISPRs was the identification of so-called *cas* genes (for CRISPR-associated) that are often found in the neighborhood of CRISPR arrays (15). Bioinformatics analyses expanded the known repertoire of *cas* genes (11, 16-19), shed light on their evolutionary origins (20, 21) and led to a comprehensive classification of *cas* gene combinations into two classes, and an increasing number of types and subtypes of CRISPR-Cas systems that differ in key mechanistic aspects (12).

The original idea that CRISPR-Cas is an adaptive immune system came from observations that sequences in CRISPR arrays on the chromosomes of bacteria match those of phage and other foreign genetic elements (10, 22, 23). *Cas* genes were – based on their domains and predicted catalytic activities – suggested to encode the protein machinery that carries out the various steps of the immune response (11). The first experimental evidence in support of this hypothesis came from a joint effort by industrial and academic partners, who showed that the lactic acid bacterium *Streptococcus thermophilus* acquired post-infection resistance (hereafter immunity) against viral infection by inserting sequences from the viral genome into CRISPR loci on the bacterial genome (24). Viruses in turn were found to overcome CRISPR-based immunity by mutation of the target sequence on their genome (25). Together, these findings fueled models of how bacteria with CRISPR systems and their viruses might coevolve (26-28). In parallel with this experimental work, genomic data suggested that CRISPR loci evolve rapidly in natural populations of acidophilic bacteria (29), and that the DNA sequences between these palindromic repeats, spacers, were homologous to that of phage, plasmids and transposons (30).

Since these pioneering studies, spacer uptake from phages and other mobile genetic elements in bacteria and archaea from natural and human-associated environments has been inferred from variation in spacer sequences within and between populations of the same species and from their homology to mobile genetic element (MGE) genomes (31-40). Experimental observations of spacer uptake in the lab in response to plasmid and phage infection have been made amongst others in engineered *E. coli* strains (41-43) and *Staphylococcus aureus* (44-47) and in wild type *Pectobacterium atrosepticum* (48), *Pseudomonas aeruginosa* (49, 50), *Roseburia intestinalis* (51), *Sulfolobus solfataricus* (52), *Streptococcus mutans* (37) and other species (reviewed in (53)). Consistent with the hypothesis that CRISPR-Cas protects bacteria from infections, some mobile genetic elements encode so-called anti-CRISPR genes (reviewed in (54)).

### How important is CRISPR to the ecology and evolution of bacteria and archaea?

While the preceding evidence and arguments demonstrate that CRISPR-Cas can protect bacteria and archaea from infectious DNAs, it is not at all clear how commonly CRISPR plays this role in natural populations of these microbes. If CRISPR commonly protects bacteria from infectious DNAs, one might expect a strong negative correlation between the presence of a CRISPR system and the signatures of horizontal gene transfer in the same genomes. For Restriction-Modification systems – the most prevalent innate immune system of bacteria and archaea – such correlations can be readily detected (55). Yet, in the case of CRISPR-Cas the evidence is ambiguous, with some studies suggesting that CRISPR does form a barrier for the movement of mobile genes between microbial species (56, 57), whereas other studies arrive at the opposite conclusion (58-60). Furthermore, most spacers from sequenced isolates have no homology to viral or plasmid sequences in databases (40) and the same holds for spacers extracted from metagenomes (39).

If indeed CRISPR-Cas is commonly important for protecting populations of microbes from phage and preventing the acquisition of mobile genetic elements, there should be no trouble detecting (in the lab or in nature) CRISPR-Cas-encoding bacteria and archaea that acquire spacers from novel sources of infectious DNAs to become immune to those infections. However, other than the bacterial species listed above, there are very few wildtype bacteria or archaea for which spacer acquisition from phage or plasmids has been demonstrated to occur at observable frequencies. Moreover, even for species that have been reported to acquire spacers, it is not clear whether they do so in response to only few or many phages and plasmids. Because in most bacteria and archaea spacer acquisition is rare, a range of elegant tricks have been developed to detect these events (44, 61-66), and while this has propelled our understanding of the mechanisms of spacer acquisition (53), and CRISPR-mediated immunity (67), it raises questions concerning the ecological importance of CRISPR-Cas immune systems. Perhaps this is because many of the domesticated bacteria and archaea that we use for research simply lost their ability to rapidly acquire spacers? Could it be that the quest to find culturable bacteria, archaea, phage and plasmids with these properties has not be adequately extensive? Or could it be that the results of these quests are commonly negative and therefore not reported? Sequence data analysis also provides a mixed picture: CRISPR loci in some species appear to evolve rapidly, e.g. refs. (29, 30, 32) and reviewed in (68), whereas others are relatively static over long periods (60, 69).

### Open questions

We hope the preceding has convinced the reader that how commonly CRISPR-Cas systems protect populations of bacteria and archaea from infections with deleterious mobile genetic elements, and the corollary, that extant CRISPR-Cas systems are maintained by selection mediated by these infectious genetic elements, are open questions. To address these issues, we explore the answers to the following four questions:

1- Under what conditions will immunity be selected for in populations with functional CRISPR-Cas systems (CRISPR^+^ bacteria and archaea) confronted with phage and plasmids?
2- Under what conditions will bacteria and archaea with functional CRISPR-Cas immune systems invade from rare upon infection with phage and plasmids, and become established in populations that lack CRISPR-Cas systems (CRISPR^-^ bacteria and archaea)?
3- What is the contribution of CRISPR immunity to the population dynamics, ecology and evolution of prokaryotes and their mobile genetic elements?
4- What are the characteristics of mobile genetic elements that lead to spacer acquisition by bacteria during an infection?

We explore the first three of these questions in two ways, (i) with heuristic, mathematical-computer simulation models of the dynamics of microbes and infectious genetic elements, and (ii) with a review of the experimental and other empirical studies that provide some answers to these questions. We separately consider the three main sources of infectious DNAs: lytic phage, temperate phage, and conjugative plasmids. For each of these we consider the <u>**invasion conditions:**</u> the conditions under which selection mediated by these infectious mobile genetic elements will (i) lead to the ascent of immune cells in populations with functional CRISPR-Cas mediated immunity, CRISPR^+^, and (ii) the establishment of CRISPR-Cas, CRISPR^+^, in populations that do not have a functional CRISPR-Cas system, CRISPR^-^. The equations for the models used and those employed for the analysis of their properties are presented in the Supplemental Material, as are the caveats and concerns about the limitations of these models and our analyses of their properties.

### I Lytic phage

#### What the models and theory tell us

We use a simple model, that captures the molecular mechanism of Type I and II CRISPR-Cas systems, which are the most abundant systems (30% and 8% of genomes, respectively (70)). In this model, there is single population of lytic phage, V, a population of bacteria that lacks a functional CRISPR-Cas system and a population that carries a functional CRISPR-Cas system, respectively CRISPR^-^ and CRISPR^+^. The CRISPR^-^ bacteria are of two possible states, phage sensitive, S, and phage resistant, SR, due to phage receptor mutations. The CRISPR^+^ bacteria can be of three states, phage sensitive not immune, C, phage resistant, CR, due to phage receptor mutations, and phage immune, CI, due to the acquisition of a phage-targeting spacer in the CRISPR array (Figure S1). We assume that resistant bacteria, SR and CR, are refractory to the phage; the phage do not adsorb to these cells. By contrast, immune bacteria, CI, can be infected by the phage, but phages do not replicate and are lost. Resistance is acquired by random mutation, and immunity through the acquisition of a spacer by infection with the phage. Both acquisition of a resistance mutation and the acquisition of spacers are stochastic processes which we model with a Monte Carlo protocol.

First, we consider the conditions under which resistance, CR, and CRISPR-Cas–mediated immunity, CI, will invade a CRISPR^+^ phage-sensitive population, C, at equilibrium with the phage. If the CRISPR^+^ population, C, is unable to generate resistant mutants, CR, but can acquire spacers, and the likelihood of acquiring a spacer upon infection with the phage is on the order of 10^−8^ or greater, there are broad conditions under which CRISPR-Cas will be selected for and immune cells CI will become established in a CRISPR^+^ population. This can be seen from the change in the mean densities of immune cells in Figure 1A. If both immunity and resistance can be generated, both will become established (Figure 1A). For examples of the associated population dynamics, see Figure S2A-C in the supplemental material. In the simulations presented in Figure 1A, upon first encounter with the phage, immunity and resistance are equally likely to be generated and therefore resistant and immune cells are equally likely to ascend to dominate the CRISPR^+^ population (we assume no costs of resistance). If immunity is 10-fold more likely to be generated than resistance, immune cells are more likely to dominate the population (Figure S2D).

**Figure 1.**
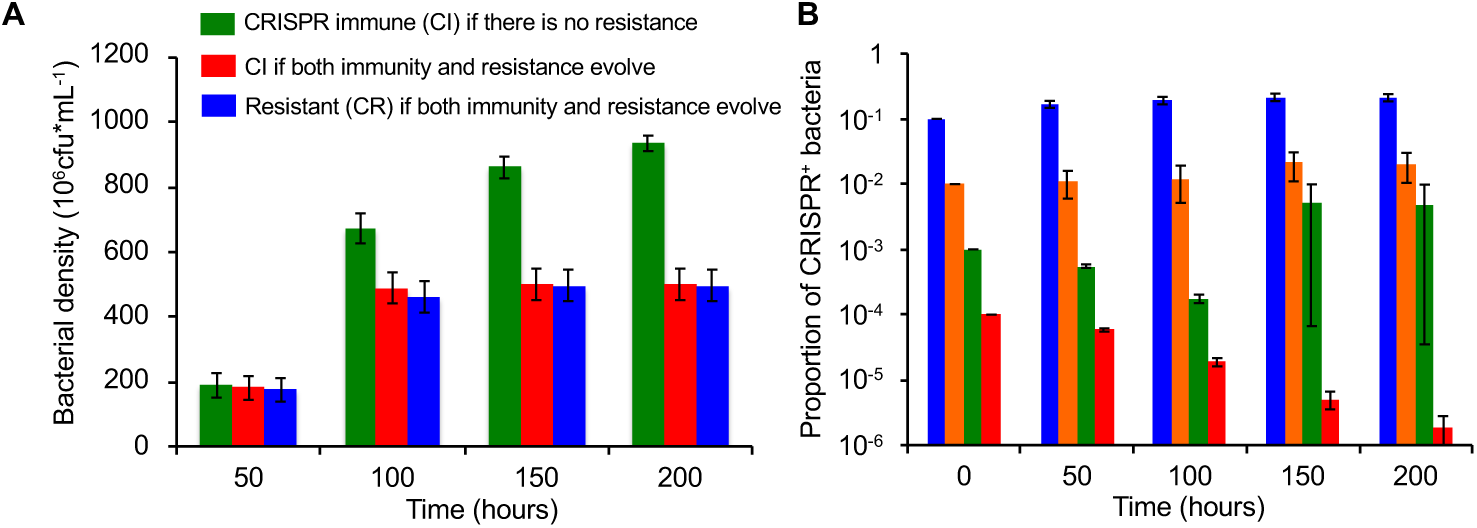
Lytic phage-mediated selection for CRISPR-Cas mediated immunity and envelope resistance. Invasion conditions: **(A)** Monte Carlo simulations of selection for CRISPR-Cas immunity and resistance in a CRISPR^+^ phage-sensitive population initially at equilibrium with the phage. Mean and standard error of the changes in density of CRISPR^+^ immune and resistant bacteria across 100 runs. Green bars are for a population of CRISPR^+^ bacteria that cannot generate resistant mutants. Red and blue bars are for populations that can evolve both CRISPR-Cas immunity (red bars) and envelope resistance (blue bars) with equal probabilities. **(B)** Invasion of CRISPR^+^ into a population of CRISPR^-^ sensitive bacteria at equilibrium with phage, with different initial frequencies of CRISPR^+^ (blue=0.1, orange=0.01, green=0.001, red=0.0001). Resistant and immune bacteria can be generated in the invading CRISPR^+^ population and resistance can be generated in the initially dominant CRISPR^-^ population. Mean and standard error of the frequency of CRISPR^+^ bacteria over time across 200 runs. Parameters: v_C_=v_CI_=v_CR_=0.7, δ=10^−7^, β=50, e=5×10^−7^, k=1, RR=500, w=0.1, r(0)=500, µ_SR_=10^−8^, µ_RS_=10^−8^, x=1.667×10^−8^ and the total volume of the vessel is Vol=100 ml. The initial densities of bacteria and phage in these simulations are at the equilibrium for a phage-limited population, respectively, C*=2×10^4^, V*=6×10^6^.

The conditions for bacteria with CRISPR-Cas (CRISPR^+^) to become established in a CRISPR^-^ population of sensitive bacteria at equilibrium with a lytic phage are more restrictive than that for CRISPR-Cas-mediated immunity to become established in a population of CRISPR^+^ bacteria. When the frequency of the invading population of CRISPR^+^ cells is low, 10^−4^ or less, the CRISPR^+^ population does not become established (Figure 1B). Moreover, in only 2 or the 200 runs did the CRISPR^+^ become invade when the initial frequency of CRISPR^+^ was 10^−3^. The reason for the difficulty of CRISPR^+^ bacteria to become established in CRISPR^-^ populations when bacteria with this immune system are initially rare is that, when confronted with phage, resistant mutants are likely to be generated in the dominant population of CRISPR^-^ sensitive cells (see Figure S3A-C). The conditions for the invasion of CRISPR^+^ in a CRISPR^-^ population are greater if the rate of spacer acquisition is higher or if the CRISPR^-^ population is unable to generate resistant mutants (Supplemental Figure S3DE)

The invasion theory considered above addresses only one element of the role of phage in the evolution of CRISPR-Cas. The other element is the length of time selection mediated by lytic phage will favor CRISPR-Cas immune systems. This will depend on (i) the capacity of the phage to generate protospacer mutants, (ii) the capacity of the bacteria to acquire novel spacers to counter protospacer mutations in the phage, (iii) the rates at which these novel spacers are acquired, (iv) the rate of mutation to resistance, and (v) the fitness costs associated with the carriage of CRISPR-Cas, surface resistance, and protospacer mutations in the phage. Some theoretical studies have partly addressed this issue (27, 28, 71), but the models employed do not consider all five of the above-listed factors contributing to the length of time selection for CRISPR-Cas will be maintained.

#### What the experiments and genomic data say

While we are unaware of experimental studies that examine the invasion of CRISPR^+^ into established CRISPR^-^ populations of bacteria or archaea, there have been several experimental studies of the population and evolutionary dynamics of lytic phage and bacteria with CRISPR-Cas systems. As anticipated by the models, CRISPR-Cas immunity readily evolves in *S. thermophilus* strains exposed to virulent phage (24, 72). In this system, bacteria with envelope resistance are normally not detected and an extended spacer – protospacer arms race can ensue when these bacteria and phage are together in serial transfer culture (24, 73-75). While the phage will eventually be lost, the duration of the arms race and the diversity of spacers and phage protospacer mutants that evolves during this process can be substantial. In these experimental populations, the densities of bacteria remain at levels similar to that of phage-free populations. Stated another way, the bacterial populations are limited by resources, rather than the phage.

In these experiments, it is clear that the co-evolutionary dynamics observed for *S. thermophilus* and its phage can be attributed to CRISPR-Cas-mediated immunity to the phage. Resistant bacteria only evolve if the CRISPR-Cas system is inactivated by either anti-sense RNA expression (76) or an antiCRISPR (*acr*) gene encoded by the phage (77), with resistance due to mutations in either the receptor or intracellular host genes required for completing the phage life cycle. This system, therefore, corresponds well with the theoretical scenario in Figure 1A (green bars), which therefore may explain why the evolution of CRISPR immunity is so commonly observed in this model organism.

Also consistent with the theoretical predictions are the results of experiments with bacteria that can evolve both CRISPR immunity by the acquisition of spacers and resistance by mutation of the phage receptors. For example, *Pseudomonas aeruginosa* strain PA14 either evolves resistance against phage DMS3*vir* (a temperate phage locked in the lytic cycle) by mutation of the Type IV pilus or immunity by the acquisition of spacers into its CRISPR arrays (49, 50). Experimental manipulation of the bacterial mutation rate shows that which of these two defense mechanisms prevails strongly depends on the rate at which receptor mutants are generated in the population (78). Similarly, the rates of spacer acquisition matters: bacteria with “primed” CRISPR-Cas systems acquire spacers at a higher rate compared to bacteria with naïve CRISPR-Cas systems; this translates in a strong increase in the proportion of bacteria that evolved CRISPR-Cas immunity following phage exposure (41, 42, 48, 61, 79-82). This phenomenon was observed first for many Type I where it relies on an imperfect match between a pre-existing spacer and the infectious genome (see ref. (53) for mechanistic details) and, more recently, a similar mechanism was also observed for Type II CRISPR-Cas systems (47, 79). When bacteria are exposed to defective phage, or when bacteria carry both a restriction-modification and a CRISPR-Cas system, the rates of spacer acquisition are also elevated, and again this leads to higher levels of evolved CRISPR immunity (83, 84). The typically low frequencies of CRISPR immunity that many bacteria evolve in the lab may therefore be at least in part explained by the high mutation rates and large population sizes relative to the rates of spacer acquisition in many model systems, although the fitness costs and benefits of CRISPR-based immunity and surface-based resistance will also be important (50, 85), which has been reviewed elsewhere (86).

### II Temperate phage

#### What the models and theory tell us: 3

In our model, (depicted in Figure S4), there is a single population of free temperate phage, P, and five populations of bacteria. Bacteria that lack a functional CRISPR-Cas system, CRISPR^-^, can exist in two states: susceptible non-lysogens, S, and lysogens, L. Bacteria that carry a functional CRISPR-Cas system, CRISPR^+^, can exist in three states: sensitive non-lysogens, C, lysogens, CL, or CRISPR-immune, CI. Infections occur at random at a rate proportional to the product of the densities of free temperate phage, P, and the bacteria. Lysogens and CRISPR immune bacteria can be infected by free phage, but the phage do not replicate and are removed from the free phage population. Upon infection of CRISPR^-^ sensitive cells, S, a fraction λ (0 ≤ λ ≤1) produce lysogens, L. The remaining (1-λ) infections results in lysis of the host to generate free phage (i.e. the lytic cycle) (87). Infections of CRISPR^+^ sensitive cells, C, have three possible outcomes: a fraction λ (0 ≤ λ ≤1-x) produce lysogens, CL with the remaining (1-x-λ) producing lytic infections and killing the infected host. By the acquisition of a spacer from the phage, a fraction x (0≤ x ≤ 1) of the infections produce CRISPR-Cas immune cells, CI. In addition, at a rate y (0 ≤ y ≤1) CRISPR-Cas immune cells are generated by lysogens, through the acquisition of a spacer from the prophage. For this analysis, we are not considering lytic mutants of the temperate phage or envelope resistant mutants.

In Supplemental Figure S5 and Figure 2 below, we consider the conditions in which selection mediated by temperate phage will lead to the establishment of CRISPR-Cas immunity in a CRISPR^+^ population and the establishment of bacteria with CRISPR-Cas (CRISPR^+^) in a CRISPR^-^ population. During the early stages of infection, temperate phages mostly transmit horizontally through the lytic cycle and selection for CRISPR immunity will therefore initially be similar to that observed for the lytic phage (Figure 1 and S5A). Following this, a large subpopulation of lysogens will form that coexists with bacteria that acquired CRISPR immunity in response to horizontally transmitting phage (88). To explore the long-term selection pressures for CRISPR immunity and invasion of CRISPR^+^ into CRISPR^-^ populations, we start our simulations when the bacterial populations are at equilibrium with the temperate phage and therefore dominated by lysogens (Figure S5B). Under those conditions, for selection mediated by temperate phage to lead to the establishment of CRISPR-Cas immune cells in populations with functional CRISPR-Cas systems, either lysogens have to generate immune cells or the prophage has to reduce the fitness of lysogens relative to non-lysogens (Figure 2A). The rate of ascent of the immune cells is inversely proportional to the fitness of the lysogens. For selection mediated by temperate phage to lead to the invasion of CRISPR^+^ bacteria in a CRISPR^-^ population, the fitness of lysogens needs to be less than or equal to that of non-lysogens. If the carriage of the prophage augments the fitness of the bacteria, as may be the case (89, 90), CRISPR immune bacteria will be selected against and the CRISPR^+^ population will not be able to invade a population without this immune system (Figure 2B). (88)

**Figure 2.**
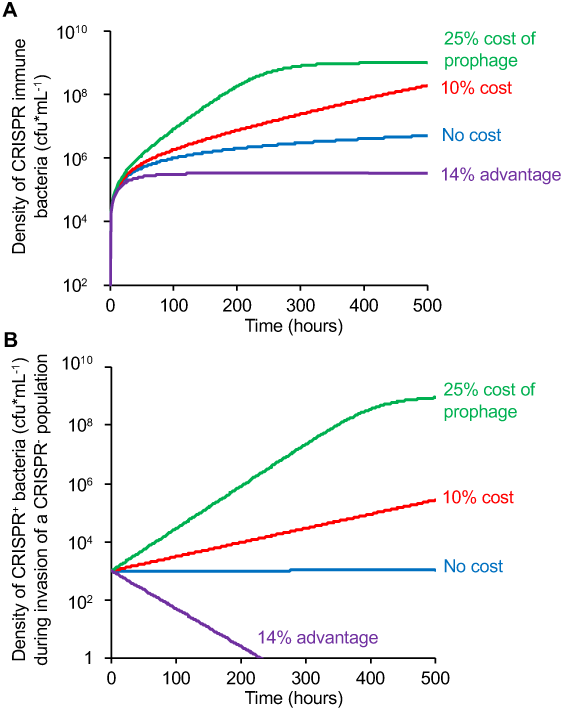
Selection-mediated by temperate phage. (A) The establishment of CRISPR immunity in a population of CRISPR^+^ bacteria at equilibrium with a temperate phage. Changes in the density of CRISPR-Cas immune bacteria are depicted. (B) The invasion of CRISPR^+^ bacteria into a population of CRISPR^-^ bacteria at equilibrium with a temperate phage. Changes in the densities of CRISPR^+^ bacteria are depicted. Carrying the prophage is associated with either a 14% advantage, a 10% cost, a 25% cost, or no cost, as indicated in the figure. For the parameter values, see Figure S5C and F.

#### What the experiments and data say

There is evidence that CRISPR-Cas systems target temperate phage in nature. For example, spacers encoded by *P. aeruginosa* isolates with Type I CRISPR-Cas systems from cystic fibrosis lungs were found to frequently target related groups of temperate phages (including DMS3). Surprisingly, however, in these patients, no spacer acquisition was observed over time (60). By contrast, in an experimental study where a wound model was infected with a mix of 6 *P. aeruginosa* strains, CRISPR immunity was found to evolve in *P. aeruginosa* strain PA14 against a prophage carried by one of the strains, known as strain B23-2 (91). Another recent study with *P. aeruginosa* PA14 and its phage DMS3 showed that carrying a primed CRISPR-Cas immune system is, in fact, maladaptive during temperate phage infection, due to immunopathology in CRISPR^+^ lysogens since the partial matching spacer triggers cleavage of the prophage. The associated fitness costs caused a rapid invasion of spontaneous mutants that had lost their CRISPR-Cas immune system (88). Acquisition of perfectly matching spacers in lysogens amplified these fitness costs, since this programs the immune system to attack the prophage inside the bacterial genome even more strongly. Such self-targeting by CRISPR-Cas is well known to be highly toxic (92-99), even for Type III CRISPR-Cas systems that target only transcriptionally active DNA (100, 101). Finally, a recent and exciting study showed that *R. intestinalis* in the mouse gut can evolve high levels of CRISPR-based immunity when one of its active prophages evolves to become hypervirulent (i.e. virulent phage mutants that can infect the lysogen) (51). This, however, brings us to what is anticipated for lytic phage. We are unaware of empirical studies that have explored the contribution of temperate phage to the establishment of CRISPR-Cas in CRISPR^-^ populations.

### III Conjugative Plasmids

#### What the models and theory tell us

In our model of the population dynamics of CRISPR-Cas immunity and conjugative plasmids there are five populations of bacteria (Figure S6). Two lack a functional CRISPR-Cas system, are CRISPR^-^, and can exist in two states: one carrying the plasmid and one not, respectively DP and S. Three populations carry a functional CRISPR-Cas system, CRISPR^+^, and exist in three states: plasmid-free, C, plasmid-bearing, CP and one that is plasmid-free but immune to the acquisition of the plasmid, CI. Plasmid transfer is by conjugation, which occurs via random contact between plasmid-bearing and plasmid-free cells (102). A “mating” between DP or CP and S produces SP. A fraction x (0 ≤ x ≤ 1) of matings between DP or CP with C produce CP, and the remaining fraction of the mating’s, (1-x), produce cells that remain plasmid-free but have acquired CRISPR immunity, CI. These CRISPR-Cas immune cells do not take up the plasmid.

The population dynamics of temperate phage and conjugative plasmids are different (87, 102) (compare the temperate phage dynamics in Figures S5A and D to the plasmid dynamics in Figures S7A and D). On the other hand, the conditions under which these infectiously transmitted genetic elements will favor the invasion of CRISPR-Cas immune bacteria into CRISPR^+^ populations and CRISPR^+^ bacteria into CRISPR^-^ populations are virtually identical (Figures S7C and F). The reason for this is that simulations are initiated at equilibrium, and with the parameters used both of these infectiously transmitted genetic elements will sweep though the bacterial populations and at equilibrium cells carrying them are the dominant population. Whether CRISPR-Cas immune cells will invade and become established in populations with lysogens and bacteria carrying conjugative plasmids depends on the fitness cost of these genetic elements: the greater the cost the broader the conditions for the establishment and maintenance of these elements. As with temperate phage, if the plasmid increases the fitness of the bacteria, CRISPR-Cas immunity would be selected against (103).

#### What the experiments and data say

Experimental studies demonstrate that bacteria can evolve CRISPR-based immunity against plasmids, and in the case of Type I CRISPR-Cas systems, spacer acquisition is accelerated if the CRISPR immune system is primed. Most commonly this priming is accomplished by engineering the plasmid in a way that it contains a sequence with a partial match to a pre-existing spacer on the genome of a CRISPR^+^ host (41, 48, 61, 104). As anticipated by the model, CRISPR immunity will be selected against if the plasmid provides a net benefit to the host, for example when it confers resistance to an antibiotic that is present in the environment. Rare mutants that lack an intact CRISPR-Cas immune system will quickly replace the dominant CRISPR^+^ population (105). The model predicts that if the carriage of the plasmid engenders a fitness cost, CRISPR-mediated immunity to that plasmid will be favored. To our knowledge, this has not been demonstrated experimentally. Indeed, the evidence we are aware of is inconsistent with this hypothesis. A study with engineered *E. coli* strains showed that even under these conditions CRISPR immunity can be maladaptive, because the time between infection and clearance of the plasmid may allow for the expression of toxin/anti-toxin cassettes. This, in turn, triggers a significant cost of plasmid removal because (short-lived) antitoxin is no longer produced to neutralize the long-lived toxin molecules (106). How common this is, is not at all clear. We anticipate that CRISPR-immunity will be favored when the plasmid engenders a fitness cost, provided that any costs of plasmid removal do not outweigh the benefits of being plasmid-free. We are unaware of any experimental or other empirical studies that have addressed the question of the conditions under which plasmid-mediated selection will favor the invasion of bacteria with CRISPR-Cas systems into CRISPR^-^ populations.

#### Other reasons why virulent phage may not select for CRISPR immunity

In the heuristic model considered here and the experiments described in the preceding section, the acquisition of spacers confers immunity to the mobile genetic element from whence the spacers are derived. This may not always be the case. More and more mobile genetic elements are found to encode anti-CRISPR (*acr*) genes that suppress CRISPR-Cas immune systems through a range of different mechanisms and with often high specificity for a single subtype (107-116). The ability of *acrs* to bypass immunity of bacteria that are already CRISPR immune varies (117, 118), but even the weakest *acr* genes characterized to date effectively block the evolution of CRISPR immunity (119).

Other viruses can by-pass CRISPR immunity without a need for *acr* genes. Some “jumbophages”, which are a class of phages with genome sizes that exceed 200kb, have been reported to form nucleus-like structure during infection (120-122). These structures contain the phage genomes, which shields it from DNA-targeting CRISPR-Cas systems, but not from systems that have RNA-targeting activity, such as Type III-A and VI-A CRISPR-Cas (123, 124). This variation in the level of protection explains why in nature spacer acquisition from nucleus-forming jumbophages is detected more frequently for bacteria that carry Type III systems compared to those that carry Type I-E or I-F systems (123). A lack of protection by CRISPR immunity is not limited to jumbophages: *E. coli* strains carrying Type I-E CRISPR-Cas that were engineered to carry a single targeting spacer against different phages revealed a lack of protection against phages R1-37 (a giant phage) and T4 (125). The ability of phage T4 to by-pass Type I-E CRISPR immunity is at least in part attributable to their genome containing glucosyl-5-hydroxymethylcytosine instead of cytosine (126), and this cytosine modification also confers infectivity to the phage when bacteria have Type II-A CRISPR-based immunity (126, 127), but not when they have Type V-A CRISPR-based immunity (126). Type I-E CRISPR-Cas offers protection against phage T7, but only under low phage densities; at high MOIs the cultures were lysed as efficiently as uninduced controls. Efficient protection against T5 was only observed if the CRISPR spacer targeted a pre-early gene (125). Furthermore, a recent study demonstrated that a Type I-F CRISPR-Cas system of *Pectobacterium atrosepticum* reduced the efficiency of plaquing of two virulent phages ϕTE and ϕM1 when the immune system was engineered to carry spacers targeting these phages. Interestingly, CRISPR-Cas was unable to rescue the host from cell death, hence resulting in an abortive infection (Abi)-like phenotype that blocks phage propagation (128). It remains to be determined if and when CRISPR immunity would evolve in bacterial populations exposed to these phages.

These observations are in stark contrast with the high levels of protection against virulent mutants of temperate phage. Examples include high levels of CRISPR-based immunity observed in *P. aeruginosa* strain PA14 against its phage DMS3vir, which we discussed above. Mild or strong overexpression of the Type I-E CRISPR-Cas system of *E. coli* targeting the non-lysogenic mutant λvir provides full immunity, with efficiency of plaquing around 10^−6^ (125, 129). Similarly, the Type II-A CRISPR-Cas immune system of *Streptococcus pyogenes* SF370 provides high levels of immunity when expressed in *S. aureus* RN4220 against the staphylococcal phage ϕNM4γ4, a lytic mutant of ϕNM4 (44-46) and the Type III-A from *S. epidermidis* RP62a provides high levels of immunity when expressed in *S. aureus* RN4220 against phage ϕNM1γ6, a lytic mutant of the temperate phage ϕNM1 (101). Mapping the variability in the levels of protection conferred by CRISPR-Cas immunity using a wider range of CRISPR immune systems and phages will be critical to understand when and where these systems matter.

### Consequences of CRISPR-Cas immunity to the population dynamics, ecology and (co)evolution of bacteria and lytic phage

In experiments where bacteria evolve high levels of CRISPR-based immunity, three possible outcomes have been observed: the phage (i) are be eliminated in short order, (ii) persist for an extended time in spacer-protospacer arms race but are eventually to be lost, (iii) persist without co-evolution. The first outcome is observed when *P. aeruginosa* PA14 is infected with DMS3*vir* (130, 131). This is because in this empirical system, the bacteria acquire many different spacers at the population level, which increases the degree of protection since it reduces the evolution and spread of phage that overcome host immunity (88, 130, 132), which they can do by point mutation (25) or deletion of the target regions on the phage genome (133). A spacer-protospacer arms race is observed when the spacer diversity at the population-level is reduced. In this case, the phage can evolve to overcome host immunity, and hosts, in turn, need to acquire novel spacers to regain immunity. This coevolutionary interaction is observed when *S. thermophilus* is infected by the lytic phage 2972: bacteria and phage coexist and coevolve for an extended time eventually for the phage to be lost (73, 75, 134). Phage extinction is in this system due to the arms-race being asymmetrical: acquisition of novel spacers is cost-free for the host (135), whereas accumulating point mutations reduces the fitness of the phage (136). Moreover, the host population gradually increases the diversity of spacers, which makes it harder for the phage to keep up with the host (75). Finally, the phage may continue to be maintained without co-evolution, when bacteria with CRISPR immunity in the population continuously acquire mutations in their CRISPR-Cas immune systems that cause phenotypic reversion to sensitivity (74), or when there is a continuous influx of sensitive bacteria due to immigration (137). While important progress has been made in understanding the consequences of the evolution of CRISPR immunity, most studies have been carried out in highly simplified environments with a single host species infected with a single phage in well-mixed and nutrient-rich broth. Future studies that examine these interactions under more ecologically relevant conditions are desperately needed to understand how CRISPR-Cas systems shape microbial population and evolutionary dynamics in nature.

## Conclusion and future directions

We do not question the validity of the hypothesis that CRISPR-Cas systems can protect bacteria and archaea from infections with deleterious mobile genetic elements. However, what remains unclear is the magnitude of the contribution of these systems to the ecology and evolution of populations of bacteria and archaea and their phage and other mobile genetic elements, collectively MGEs. As outlined above, many key questions remain. First, more experimental and observational studies are needed to understand not only how frequently, but also when, where and why CRISPR-Cas systems play a role in defense against MGEs (138). Second, it remains unclear how commonly selection mediated by MGEs is responsible for the existence and maintenance of CRISPR-Cas systems in populations of bacteria and archaea, and how this is determined by the type of MGE. Third, while CRISPR-Cas systems clearly spread by HGT, it remains unclear how these genes are able to invade a population from rare, especially if the bacteria or archaea can evolve envelope resistance as well. Finally, our understanding of the ecological and evolutionary consequences of CRISPR-Cas immune responses are limited to *in vitro* experiments that lack much of the biotic and abiotic complexity of natural environments. Could it be that the biotic and abiotic complexity of the real world, where communities of microbes include multiple species and strains as well as diversity in phage and plasmids, are spatially structured and exist in fluctuating environments are critical to the evolution and maintenance of CRISPR (50, 85, 139, 140)? Filling these gaps in our current understanding of CRISPR ecology and evolution requires interdisciplinary approaches that combine observational studies, mathematical and computer simulation models, as well as population and evolutionary dynamics experiments. The question is how can we do experiments in a way that they would also provide a test of the generality of the hypotheses that are being examined? For that, we would need a diverse array of culturable bacteria and archaea with functional CRISPR-Cas systems and a diverse set of phage and plasmids, which inevitably requires many different research teams to examine these questions. This brings us back to our concern about the dearth of bacteria and archaea phage and plasmid systems amenable for these experimental studies, and the “fishing expedition” dilemma that a quest to find new systems engenders. However, it is always more difficult publishing negative evidence, no matter how informative that evidence would be. We argue that there is a pressing need to publish any negative results of spacer acquisition in response to MGEs, since knowing which culturable bacteria and archaea with functional CRISPR-Cas do and do not acquire spacers and how this depends on the type of infectious DNA will be critical to fully understand the evolutionary ecology of CRISPR-Cas.

## Acknowledgments

ERW received funding from the Natural Environment Research Council (http://www.nerc.ac.uk) (NE/M018350/1), the Biotechnology and Biological Sciences Research Council (BB/N017412/1), and the European Research Council (https://erc.europa.eu) (ERC-STG-2016-714478 - EVOIMMECH). BL was funded by grants from the U.S. National Institutes of General Medical Science GM091875-17 and 1R35 GM136407-01 to (BRL) and the generosity of Emory University.

## Supplemental Material. A theoretical consideration of the population and evolutionary dynamics of CRISPR-Cas mediated immunity

As considered in the body of this perspective, CRISPR-Cas can provide bacteria protection against infections by mobile genetic elements of three major sources, lytic (virulent) phage, temperate phage, and self-transmissible plasmids. In this supplemental material, we use simple mathematical and computer simulation models to elucidate the *a priori* conditions under which these three types infectious mobile genetic elements will select for (i) CRISPR-Cas mediated immunity in populations with functional CRISPR-Cas systems, CRISPR^+^, and (ii) the ascent of CRISPR^+^ bacteria in populations without functional CRISPR-Cas systems, CRISPR^-^. A particular focus of our analysis of the properties of these models are “invasion conditions”; the conditions under which selection mediated by mobile genetic elements will lead to increases in the frequency of CRISPR-Cas immune populations when they are initially rare. As in the body of this report, we separately consider these invasion conditions for populations with functional CRISPR-Cas systems, CRISPR^+^ and populations without these adaptive immune systems, CRISPR^-^.

For all of these models, we assume the populations are maintained in continuous (chemostat) culture. A limiting resource, r µg/ml from a reservoir where it is present at a concentration R µg/ml enters a habitat of unit volume at a rate w per hour, which is the same rate at which excess resources, bacteria, and free phage are removed. The rate of growth of the bacteria of type i is directly proportional to its maximum growth v_i_ per cell per hour and hyperbolic function of the concentration of the resource (141).

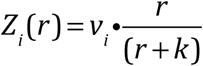

The parameter, *k*, the Monod constant, is the concentration of the limiting resource, where the rate of growth of the population is half its maximum value. As in (142), we assume the limiting resource is consumed at a rate proportional to the growth rate of those bacteria, *Z*_*i*_*(r)*, and the amount of resource required to produce a new cell is e µg/ml.

The properties of these models were analyzed numerically with Berkeley Madonna. For copies of these programs and instructions for their use, write to. The models used for these numerical analyses are generic and chosen to illustrate the conditions under which selection mediated by phage and plasmids will favor the evolution of CRISPR-Cas immunity. However, the growth rates, phage infection and lysogeny rates, and plasmid transfer rates are in the ranges estimated in the cited experimental studies.

### I. The population dynamics of lytic phage and bacteria with and without CRISPR-Cas immunity

#### The Model

In this model, depicted in Figure S1, there is a single population of lytic phage, V, and two types of bacteria: those that are CRISPR^+^ (i.e. carry a functional CRISPR-Cas system) and those that are CRISPR^-^ (i.e. lack a functional CRISPR-Cas system). CRISPR^-^ bacteria can exist in two states; they can either be sensitive to the phage or they can have surface resistance, respectively S and SR. CRISPR^+^ populations can exist in three states: one sensitive to the phage, C, one with surface resistance, CR, and one that is CRISPR immune, CI. The resistant cells, SR and CR, are refractory to the phage, while the CRISPR immune population can be infected by the phage, but the infecting phage are lost. The variables V, S, SR, C, CR, and CI are both the designations and densities of these populations, particles, in cells per ml. In Table S1, we list the variables and parameters of this model, their dimensions and the range of parameter values used.

**Figure: S1.**
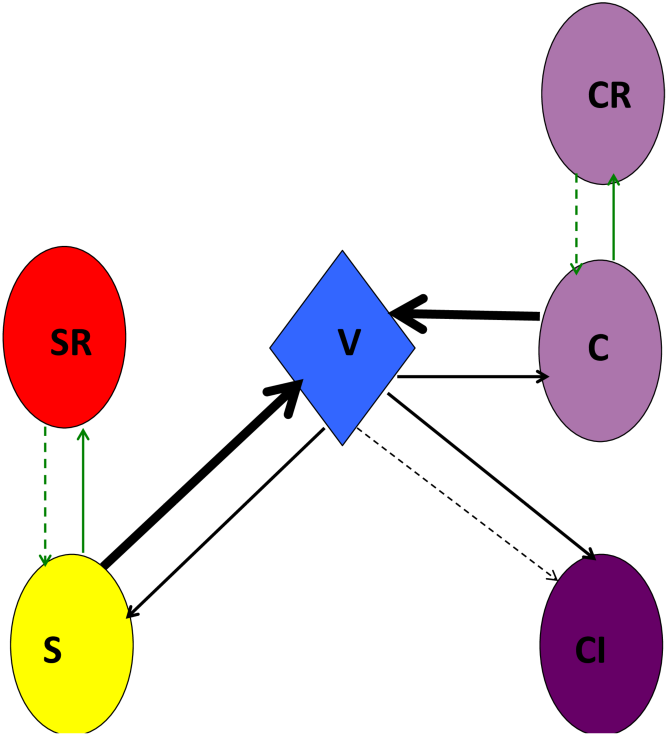
Lytic phage model: V - free phage, S - CRISPR-sensitive bacteria, SR - CRISPR-resistant (refractory) bacteria, C - CRISPR+ phage sensitive bacteria, CR - CRISPR+ resistant (refractory) bacteria, CI - CRISPR+ immune bacteria. The thin black arrows indicate infection and thick black arrow lytic production of phage. The thin green solid and dotted lines designate transitions between states due to mutation or phenotypic processes, respectively, the generation and loss of resistance and CRISPR-Cas mediated immunity.

In the absence of phage, bacteria grow at maximum rates, v_S_, v_SR_, v_C_, v_R_ and v_CI_ per cell per hour. The phage adsorb to the bacteria at a rate proportional to the product of their densities and a rate parameter, δ cell x ml/hour (143). If the hosts are sensitive to the phage, S or C, upon infection β phage particles per cell, the burst size, are produced and the infected host cells are removed from the population. Phage infecting CRISPR^+^ immune cells, CI, are lost and removed from the population. A fraction, x (0 ≤x≤ 1) of the adsorptions of phage V to sensitive CRISPR^+^ cells, C, produce immune cells CI (71). By mutation at a rate µ_SR_ per cell per hour, sensitive cells generate resistant mutants, S→SR, and C→CR. As in (144) we assume that either by mutation or phenotypic processes, at a rate µ_RS_ per cell per hour resistant cells become susceptible, SR→S, and CR→C. With these definitions, the rates of change in the densities of the bacteria and phage and concentration of resources is given by the following set of coupled differential equations.

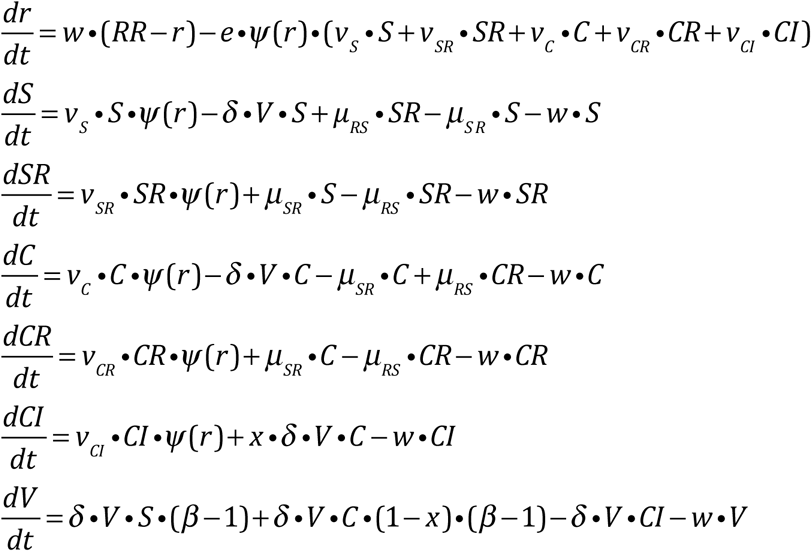

*where* 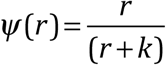

#### 1 The evolution of CRISPR immunity and surface resistance in CRISPR^+^ populations

When sensitive populations of CRISPR^-^ bacteria, S, are confronted with phage, resistant mutants, SR, will ascend and become the dominant bacterial population. In CRISPR^+^ populations this dominant population will be either resistant bacteria, CR, or immune cells, CI, which are produced by the acquisition of a spacer from the phage. Whether resistant mutants or CRISPR-Cas immune cells will dominate depends on the rate of mutation to resistance, S→SR and C→CR, and the rate of spacer acquisition, C→CI.

To illustrate this, we consider a continuous culture community of sensitive bacteria of density N cells per ml at equilibrium with lytic phage in a habitat where resources are sufficient for the rates of growth to be at their maximum. Under these conditions, the densities of the phage and bacteria populations would be,

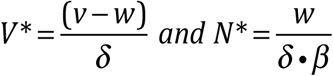

Where *V\** is the equilibrium density of the phage and *N\** the equilibrium density of the bacteria. For example, if the maximum growth rate of the bacteria, v=0.7 per hour, the adsorption rate constant, δ =10^−7^, the burst size β=50, and the flow rate w=0.1 per hour, the equilibrium densities of bacteria and phage would be, respectively N*=2.0 × 10^4^ and

V*=6.0 × 10^6^.

For a population of CRISPR^+^ sensitive bacteria at equilibrium with phage, C* and V*, whether resistant, CR, or immune CI cells will appear first will depend on the rate of mutation to resistance, µ_SR_, the likelihood of the bacteria picking up a spacer, the parameters δ and x and the total number of bacteria, C*•Vol. If µ_SR_•C*•Vol > x•δ •V*•C*•Vol, resistance will more likely appear beforeCRISPR immunity. In general, for CRISPR-Cas immune cells to be generated before resistant ones,

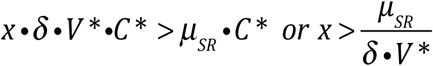

For example, if the mutation rate to resistance is µ_SR_=10^−8^, with δ=10^−7^ and V*=6×10^6^, for CI to appear before CR, the probability of acquiring a spacer upon infection, x has to exceed 1.67 × 10^−8^. If x=1.67 × 10^−7^, the probability of acquiring a spacer is 10X as great as that of generating a resistant mutant.

#### (i) Establishment of resistance and immunity in a CRISPR+ population

In Figure S2, we consider the establishment of immune and resistant bacteria, respectively CI and CR, in a phage sensitive CRISPR^+^ population, C, initially at equilibrium with the phage. For these simulations, we use a semi-stochastic version of this model where the generation of mutants or the acquisition of spacers are determined by a Monte Carlo process, with the rest of the transitions between states being deterministic. Consequently, although all of the runs were started with the same conditions, the dynamics differed because mutation and the acquisition of spacers are stochastic processes. In Figure S2A, B, and C we present runs with different outcomes: A where the immune cells become the dominant population, B where the resistant cells become the dominant population, and C where the resistant cells become the dominant population but a high density of immune cells are maintained.

For a more comprehensive perspective of when these different outcomes are anticipated and the likelihood of their occurrence, we use a Monte Carlo simulation to determine the average density of immune and resistant cells at different times for 100 runs with these parameters (Figures 1A and S2D). In the absence of mutation to resistance, µ_SR_=µ_RS_=0, within short order immune cells, CI, evolve and dominate the CRISPR^+^ population. When resistance and immune cells are equally likely to be generated, both are equally likely evolve to ascend to dominate the bacterial population. If, as in Figure S2D, CRISPR-Cas immunity is more likely to be generated than envelope resistance, immune cells evolve to dominate.

#### (ii) Establishment of CRISPR^+^ bacteria in CRISPR^-^ populations

In Figure S3A-C, we consider the dynamics of the changes in the densities of the different populations of bacteria, S, SR, C, CI and CR, the phage V, and the concentration of the resource, r, for situations where resistance can be generated µ_SR_=µ_RS_=10^−8^. In these simulations, the sensitive populations are initially at equilibrium with the phage, V= 6×10^6^, and the density of CRISPR^+^ is around 10% of the total population, respectively C=2×10^3^ and S=1.8×10^4^ cells ml^-1^. CRISPR^-^, SR, or CRISPR^+^, CR, cells can emerge and ascend to dominate the population (Figure S3A and B, respectively). CRISPR^+^ immune cells, CI, can also ascend to dominate the bacterial population (Figure S3C). In this latter simulation the probability of the acquisition of spacer upon infection was 10 times as great as that in Figures S3A and B, x=1.667x 10^−7^ rather than x=1.667x 10^−8^.

**Figure S2:**
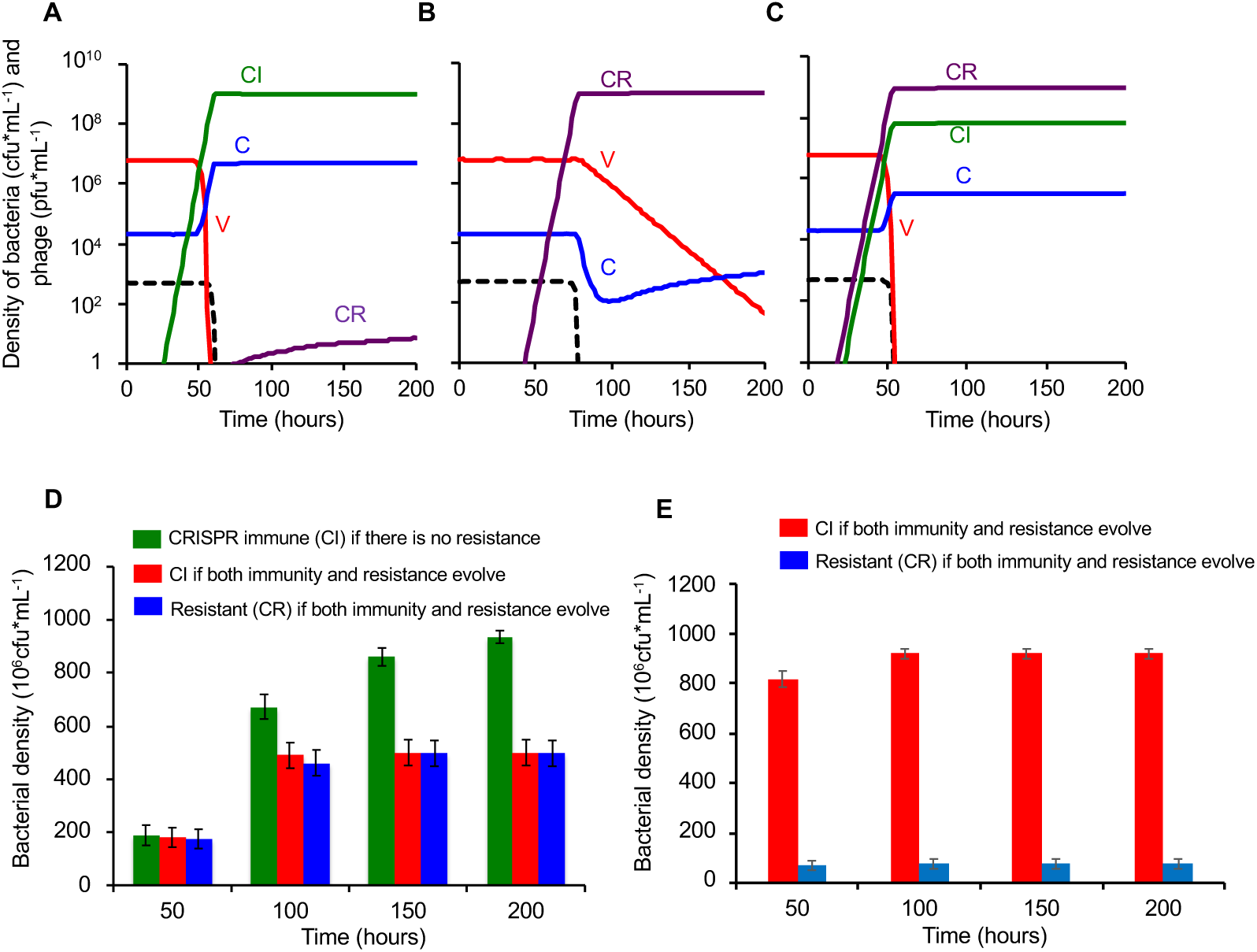
Establishment of immune, CI, and resistant, CR, cells in a sensitive CRISPR^+^ population, C. Standard parameters v_C_=v_CI_=v_CR_= 0.7, δ=10^−7^, β=50, e=5×10^−7^, k=1, RR=500, w=0.1, r(0) =500, µ_SR_=10^−8^ µ_RS_=10^−8^, and the total volume of the vessel is Vol=100 ml. The initial densities of bacteria and phage in these simulations are at the equilibrium for a phage-limited population, respectively, C^*^ =2×10^4^, V^*^ =6×10^6^, and x=1.667×10^−8^ unless indicated otherwise. Changes in the densities of bacteria and phage, when A) immunity, CI, arises and ascends to dominance before resistance, CR; B) resistance, CR, emerges and ascends to dominance before immune cells are generated or C) resistance emerges and ascends shortly before immune cells emerge. D) The outcome of 100 independent simulations, mean and standard error of the density of CI and CR at different times, where the probability of spacer acquisition x=1.667×10^−8^ (this figure is also shown in the main text, as Figure 1A) or E) or where the probability of picking up a spacers is 10X that of generating a mutant x=1.667×10^−7^. Black dotted lines are resource concentration.

To determine the relationship between the initial frequencies of CRISPR^+^ and CRISPR^-^ on the conditions for CRISPR^+^ cells to become established in CRISPR^-^ population, we use Monte Carlo simulations, 200 runs with each set of parameters. We follow the changes in the density of CRISPR^+^ cells at different at times for different initial frequencies of CRISPR^+^ cells. As our criterion for the establishment of CRISPR^+^ in a CRISPR^-^ population, we consider the change in the density of CRISPR^+^ between time 0 and 200 hours. By this criterion CRISPR^+^ can become established in populations of CRISPR^-^ when the initial frequency of CRISPR^+^ is as low as 10^−3^, depending on whether immunity is generated at a low or high rate relative to mutation (Figures S3D and S3E). In considering this, it is worth noting that in the runs with the initial frequency of CRISPR^+^ of 10^−3^, CRISPR^+^ dominated at 200 hours in only 2 out of the 200 runs, with both the lower and higher rates of CRISPR-Cas mediated immunity.

Overall, we interpret these simulation results to suggest that if the frequency of CRISPR^+^ cells that are initially neither resistant or immune to the phage is less than 10^−3^, the likelihood of CRISPR-Cas immunity successfully becoming established in a CRISPR^-^ population at equilibrium with the phage is negligible. This is particularly important when considering the establishment of CRISPR^+^ cells in CRISPR^-^ populations by horizontal gene transfer when the initial frequency of CRISPR^+^ cells is going to be low.

**Figure S3.**
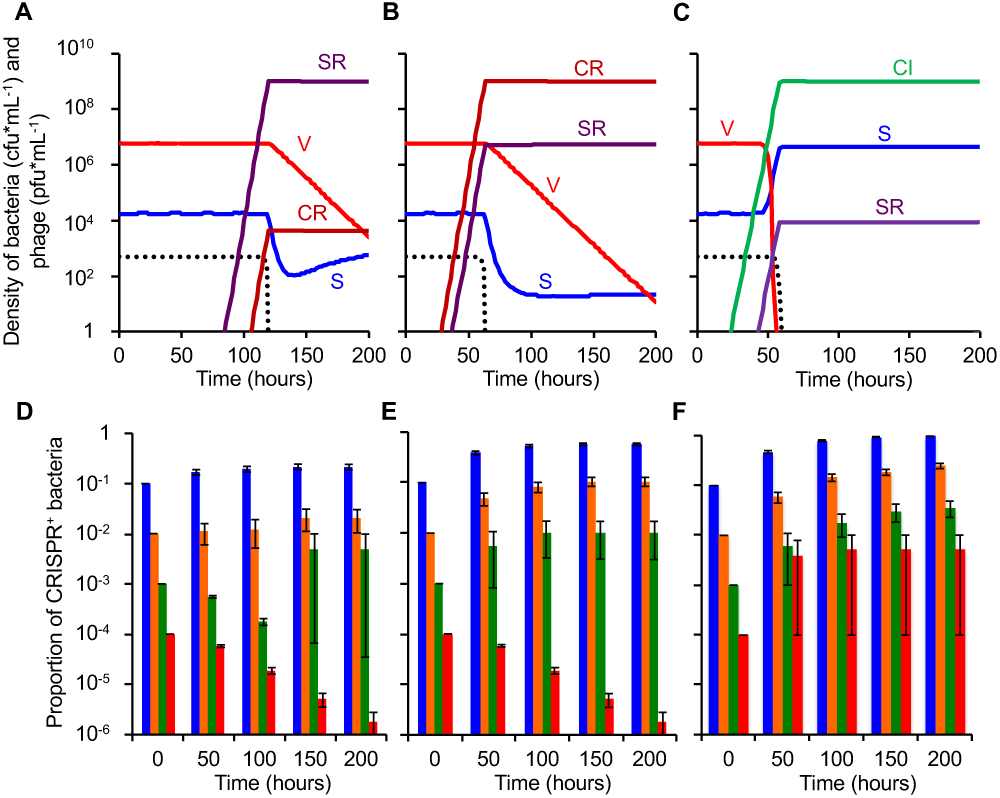
Establishment of CRISPR^+^ in a CRISPR^-^ phage sensitive population at equilibrium with lytic phage. Standard parameters, v_S_=v_R_=v_C_=v_CI_=v_CR_= 0.7, δ=10^−7^, β=50, e=5×10^−7^, k=1, RR=500, w=0.1, µ_SR_=10^−8^, µ_RS_=10^−8^ (save for F) and the total volume of the vessel is Vol=100 ml. In this simulation the equilibrium population of sensitive bacteria includes both CRISPR^-^ and CRISPR^+^. A and B) changes in the densities of bacteria and phage in populations initiated with 10% CRISPR^+^, C, and 90% CRISPR^-^, S, and x=1.667×10^−8^. A) CRISPR^-^ resistant cells, SR, evolved to dominate. B) CRISPR^+^ resistant cells, CR, evolved to dominate. C) x=1.667×10^−7^, and CRISPR^-^ immune cells, CI, evolved to dominate. D) Monte Carlo simulations, mean and standard errors in the frequency of CRISPR^+^ cells in 200 simulated populations initiated with different frequencies of CRISPR^+^, C, and CRISPR^-^, S, at equilibrium with the phage with x= 1.667×10^−8^ (this figure is also shown in the main text, as Figure 1B). Blue=0.1, orange=0.01, green=0.001, red=0.0001 initial frequency of CRISPR^+^. E) Monte Carlo simulations, mean and standard errors in the frequency of CRISPR^+^ cells. 200 simulated populations initiated with different frequencies of CRISPR^+^, C, and CRISPR^-^, S, at equilibrium with the phage with x=1.667×10^−7^. Color coding as in D. F) Monte Carlo simulations, mean and standard error in the frequency of CRISPR^+^ cells. 200 simulated populations initiated with different frequencies of CRISPR^+^, C, and CRISPR^-^, S, at equilibrium with the phage with x=1.667×10^−^ 7, and no resistance possible µ_SR_=µ_RS_=0. Color coding as in D. Black dotted lines are resource concentration.

#### (iii) Caveats

In choosing the parameters for these simulations of lytic phage selecting for CRISPR-Cas mediated immunity, we are making two assumptions that can affect the predictions. One, all of the populations are equally fit (i.e. they have the same maximum growth rates). For example, if CRISPR-Cas engenders a cost relative to sensitive cells, the conditions for the establishment of immunity in a CRISPR^-^ or CRISPR^+^ population are going to be less than that predicted by this model. If CRISPR-Cas is less costly than envelope resistance, v_CI_ > v_CR_, immunity is more likely to evolve than resistance. Two, the populations of sensitive cells being invaded, C or S, are at equilibrium with the phage. If indeed these populations were confronted by phage, they may already be dominated by resistant cells.

### ii) Population dynamics of temperate phage and bacteria with and without a CRISPR-Cas system

#### The Model

In this model, diagrammed in Figure S4, there is a single population of temperate phage, P, and two types of bacteria: those that are CRISPR^+^ (i.e., carry a functional CRISPR-Cas system) and those that are CRISPR^-^ (i.e. lack a functional CRISPR-Cas system). The CRISPR^-^ bacteria can exist in two states; they can either be sensitive non-lysogens, or they can be lysogens (carry the prophage), respectively S and L. The CRISPR^+^ populations can be present in three states: sensitive non-lysogens, C, lysogens, CL, and CRISPR immune, CI. The phage can exist in three states, as free phage, P, or as prophage in CRISPR^-^ lysogens, L, or as CRISPR^+^ lysogens, CL. The bacteria grow at maximum rates, v_S_, v_L_, v_C_, v_CL_, and v_CI_ per cell per hour. The phage adsorb to the bacteria at a rate proportional to the product of their density, that of the bacteria, and a rate parameter, δ cell x ml/hour (143). The S and C populations support the lytic replication of the phage. As in (145) with a probability λ (0 ≤ λ≤1) upon infection with lytic phage, the S and C populations become lysogens, respectively L and CL. These lysogens are immune to super-infection with the temperate phage, as are the CRISPR^+^ immune cells, CI; phage that infect these immune cells are removed from the population. CRISPR^+^ immune cells, CI, are generated in two ways, from existing lysogens, at rate y per cell per hour, and by infection with P, with a probability x per infected cell. In addition to being produced by lytic infections, with a burst size β, free temperate phage are generated by induction of the lysogens, at a rate i per cell per hour. When they lose the prophage, CRISPR^-^ lysogens revert to sensitivity, S, and when they lose the prophage, CRISPR^+^ lysogens revert to sensitivity, C. In this model, we assume CRISPR-Cas is not lost.

**Figure S4.**
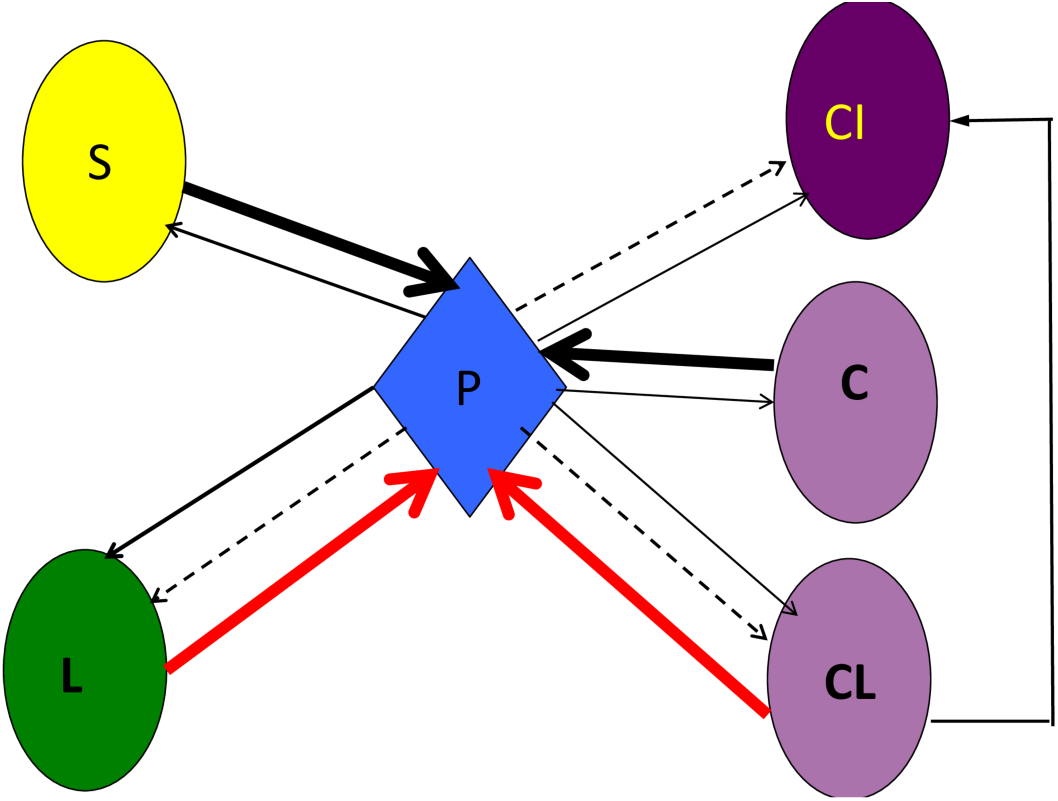
Model of the population dynamics of temperate phage and bacteria with and without CRISPR-Cas systems. P is the density of free phage, S, and L are respectively CRISPR^-^ sensitive and lysogenic bacteria. C, CL, and CI are, respectively, the designations and densities of CRISPR^+^ bacteria that are sensitive to the free phage, lysogens, and immune cells. Solid red lines denote the production of free phage by induction of the lysogens. The thick solid black lines denote the production of phage by lytic infection. The thin broken lines denote the loss of the phage due to the adsorption to immune cells, L, CL, and CI. The spacers responsible for the immunity of non-lysogens, C, are picked up by infection with the phage and from CRISPR^+^ lysogen.

With these definitions and assumptions, the rates of change in the densities of the bacterial populations, free phage, and the concentration of the limiting resource are given by the following set of coupled differential equations.

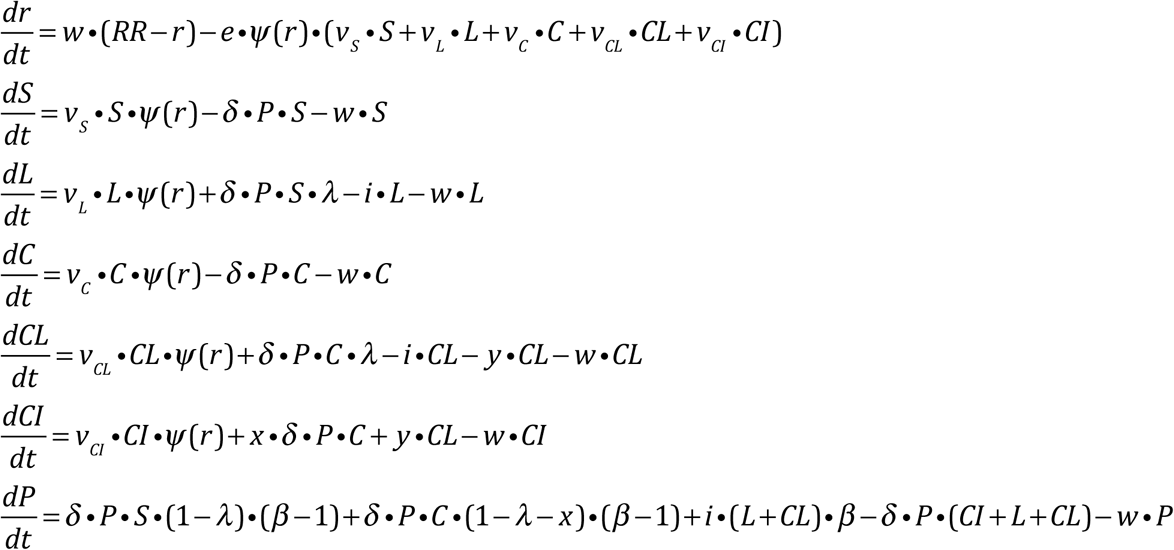

*where* 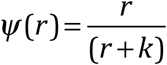

#### Temperate phage-mediated selection for CRISPR-Cas

We open our analysis of the properties of this model with an exploration of the conditions under which CRISPR-Cas immune cells, CI, will invade and become established in a CRISPR^+^ population. In Figure S5A, we consider the dynamics of the invasion of a population of bacteria immune to the temperate phage, CI, into a population initially composed of sensitive non-lysogens and free temperate phage, C and P. Due to lytic infection and the generation of lysogens, the sensitive, C population is lost. The density of the free temperate phage increases rapidly and the C population is converted into lysogens CL. In this simulation, the carriage of the prophage reduces the fitness of the lysogens, CL relative to C and CI. Immune cells, CI, are produced and, because of their fitness advantage over the lysogens, replace the lysogens as the dominant population of bacteria. The density of free phage declines when immune population ascends to dominate. In the absence of CRISPR-Cas mediated immunity, there would be a stable equilibrium with lysogens and temperate phage, CL and P (145), which can be seen in Figure S5B. In this and the following simulation, Figure S5C, we consider the invasion of CRISPR-Cas immunity, CI, in populations at equilibrium with the temperate phage. In the absence of selection for or against lysogens, the immune cells, CI, increase slowly, due to the conversion of lysogens into immune cells, CL→CI. The rate of increase of the CRISPR-Cas immune cells is proportional to the fitness cost of carrying the prophage (Figure S5C).

In Figures S5D-F, we consider the invasion of CRISPR^+^ cells in a CRISPR^-^ population. In Figure 5D we follow the dynamics of temperate phage, lysogeny and the invasion of CRISPR^+^ cells (CI and CL) in a population initiated with sensitive non-lysogens, S and temperate phage, P and a 100 CRISPR^+^ sensitive non-lysogens. The phage density increases initially and, within short order, the CRISPR^-^ lysogens, L, ascend. While the invading population of CRISPR^+^ sensitive non-lysogens, C are lost due to the phage, they are converted into lysogens, CL, which are maintained for a while as a minority population. CRISPR-Cas immune cells, CI are generated and, because in this simulation they have a selective advantage over the lysogens, they increase in density, invade, and will eventually become the dominant population of bacteria. In Figure S5E we consider a population of CRISPR^-^ lysogens at equilibrium with the temperate phage and a minority population of CRISPR immune cells, CI in the absence of selection for or against the lysogens, L and CL. Under these conditions, the density of the invading CRISPR^+^ population does not increase or decrease. If, the prophage reduces the fitness of lysogens, the CRISPR^+^ population will invade, Figure S5F with the rate of ascent inversely proportional to the costs. If the prophage augments the fitness of the lysogens, relative to non-lysogens, the CRISPR^+^ population will be selected against.

**Figure S5.**
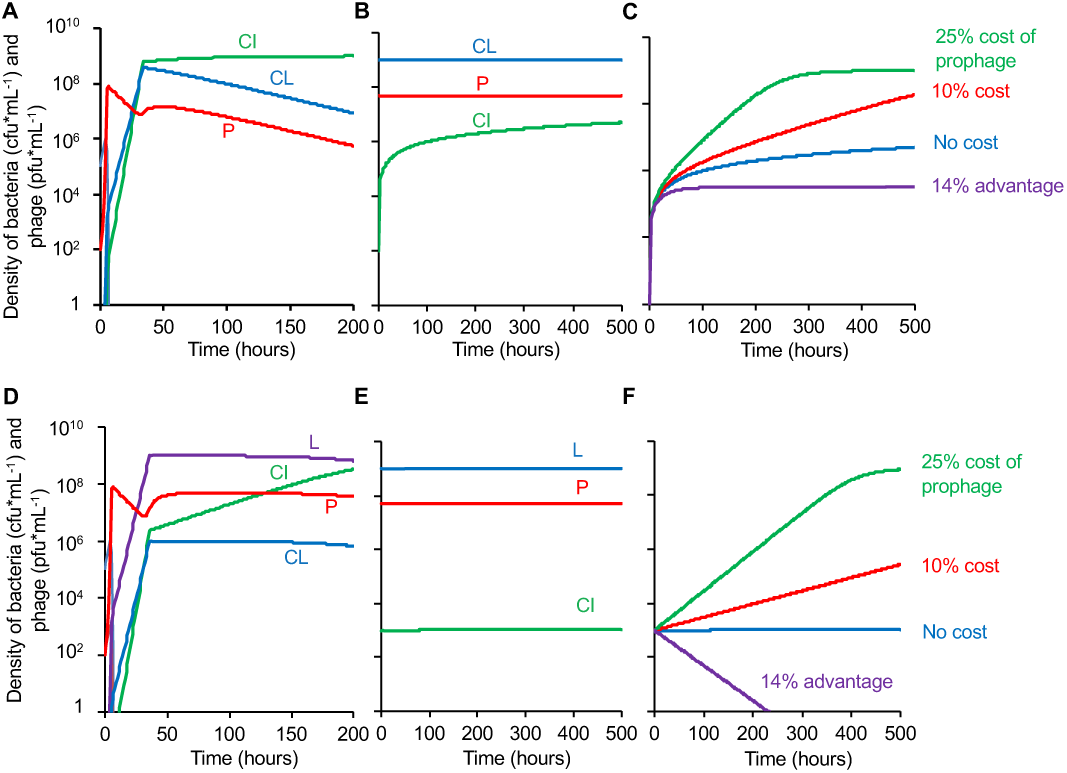
Invasion conditions for CRISPR-Cas mediated immunity in populations of bacteria and temperate phage. Standard parameters: RR=500, w=0.1, k=1.0, e=5×10^−7^, 1=10^−7^, 1=50, i=10^−4^, 1=10-3, x=y=10^−5^. A, B and C) Establishment of immunity in a CRISPR^+^ population. A) Dynamics of the changes in the densities of free phage, CRISPR^+^ lysogens, and CRISPR^+^ immune bacteria, in populations initiated with free phage and sensitive CRISPR^+^ bacteria, with a 25% fitness cost of lysogens, v_C_=v_CI_=0.70, v_CL_=0.525. The light blue line corresponds to the initially sensitive cells, which are quickly lost from the population. B and C) Populations at equilibrium with lysogens and free temperate phage. B) Changes in the densities of bacteria and phage in the absence of selection for or against lysogens, v_C_=v_CL_=v_CI_=0.7. C) Changes in the densities of CRISPR-Cas immune cells with different cost of lysogeny, no cost, v_C_=v_CI_=0.7, v_CL_=0.7, a 25% costs v_CL_=0.525, a 10% cost v_CL_=0.63, with lysogens 14.3% more fit than non-lysogens v_CL_=1.0. D, E, and F) Establishment of bacteria with CRISPR-Cas (CRISPR^+^) in a CRISPR^-^ population. D) Dynamics of the changes in density of free phage, CRISPR^-^ and CRISPR^+^ bacteria in a population initiated with free temperate phage, and CRISPR^-^ and CRISPR^+^ sensitive non-lysogens, with a 25% cost due to the carriage of the prophage v_S_=v_C_=v_CI_=0.7, v_L_=v_CL_=0.525. The light blue line corresponds to the initially sensitive cells, which are quickly lost from the population. E and F) Population dynamics of CRISPR^+^ in a population of CRISPR^-^ bacteria initially at equilibrium with the temperate phage. E) Changes in the densities of bacteria and phage in the absence of selection for or against lysogens v_S_=v_L_=v_C_=v_CL_=v_CI_=0.7, F) Changes in the densities of CRISPR^+^ cells with different costs of lysogeny, no cost v_S_=v_C_=v_CI_=0.7, v_L_=v_CL_=0.7, a 25% cost, v_L_=v_CL_=0.525, a 10% cost, v_L_=v_CL_=0.63, a 14.3% advantage v_L_=v_CL_=1.0.

In Figures S5D-F, we consider the invasion of CRISPR^+^ cells in a CRISPR^-^ population. In Figure 5D we follow the dynamics of temperate phage, lysogeny and the invasion of CRISPR^+^ cells (CI and CL) in a population initiated with sensitive non-lysogens, S and temperate phage, P and a 100 CRISPR^+^ sensitive non-lysogens. The phage density increases initially and, within short order, the CRISPR^-^ lysogens, L, ascend. While the invading population of CRISPR^+^ sensitive non-lysogens, C are lost due to the phage, they are converted into lysogens, CL, which are maintained for a while as a minority population. CRISPR-Cas immune cells, CI are generated and, because in this simulation they have a selective advantage over the lysogens, they increase in density, invade, and will eventually become the dominant population of bacteria. In Figure S5E we consider a population of CRISPR^-^ lysogens at equilibrium with the temperate phage and a minority population of CRISPR immune cells, CI in the absence of selection for or against the lysogens, L and CL. Under these conditions, the density of the invading CRISPR^+^ population does not increase or decrease. If, the prophage reduces the fitness of lysogens, the CRISPR^+^ population will invade, Figure S5F with the rate of ascent inversely proportional to the costs. If the prophage augments the fitness of the lysogens, relative to non-lysogens, the CRISPR^+^ population will be selected against.

### III Population dynamics of conjugative plasmids and bacteria with and without CRISPR-Cas mediated immunity

#### The Model

There are two populations of CRISPR^-^ cells, one that carries the plasmid and one that does not, DP, and S, respectively, and three populations of CRISPR^+^ cells, plasmid-free, plasmid-bearing, and immune, respectively C, CP, and CI. These populations grow at maximum rates, v_S_, v_P_, v_C_, v_CP_, v_CI_ per cell per respectively. The plasmids are transferred at rates proportional to the product of the densities of plasmid-bearing and plasmid-free cells and a rate parameter, γ (146). CRISPR^+^ cells, C, acquire spacers and become immune to infection with the plasmid, CI, at a rate proportional to the product of their densities, the rate constant of plasmid transfer, γ, and the probability of picking up a spacer x (0 ≤ x ≤1) upon conjugation. Immune CRISPR^+^ cells can also be generated from plasmid-bearing CP at a rate y per cell per hour. Plasmids are lost by vegetative segregation at a rate τ per cell per hour, resulting in DP cells reverting back to S and CP cells reverting back to C.

**Figure S6.**
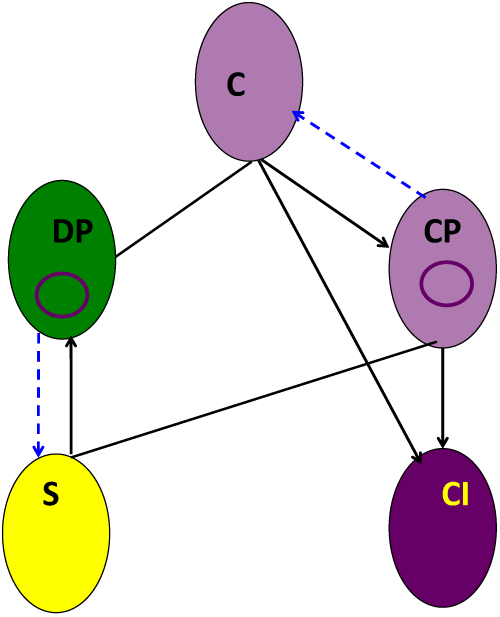
Model of the population dynamics conjugative plasmids and bacteria with and without CRISPR. There are two populations of CRISPR^-^ cells, plasmid-free, and plasmid bearing, S, and DP. There are three populations of CRISPR^+^ bacteria, those that are plasmid-free, those that carry the plasmid and those that are immune to the plasmid, respectively, C, CP, and CI. Plasmids are transferred to plasmid-free cells at a rate proportional to the product of their densities and a rate constant, γ. Immune, cells, CI are produced by infection of C by mating with a plasmid bearing cells or from a transition from CP to CI.

With these definitions and assumptions, a chemostat habitat and resource-limited growth, the rates of change in the densities of the different populations and concentration of the resource are given by the following array of coupled differential equations.

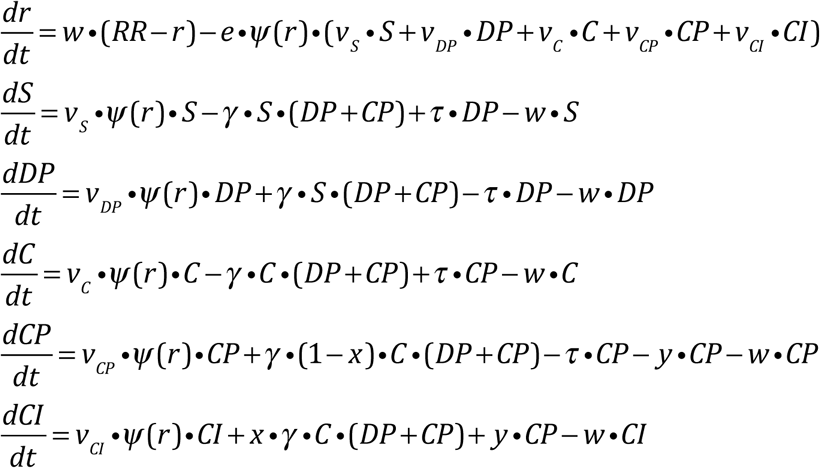

*where* 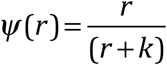

When bacteria carrying conjugative plasmids are introduced into receptive populations of plasmid-free cells in continuous culture, the plasmid can sweep through the population and convert the plasmid-free cells into plasmid bearing (146). These dynamics can be seen in Figure S7A, where we consider the invasion of a CRISPR-Cas immune population, CI, into a population of plasmid-bearing and plasmid-free CRISPR^+^ cells, C and CP. In these simulations, the plasmid engenders of 25% fitness cost; the immune bacteria, CI ascend and eventually replace the plasmid-bearing cells as the dominant population. With the parameters considered in the simulations, in the absence of immunity, CI, there are stable equilibria between the plasmid-bearing and plasmid-free bacteria (146). In the absence of a cost due to the carriage of the plasmid, the initially rare immune population, CI, increase in density in a CRISPR^+^ population at equilibrium with the plasmid (Figure S7B). This increase in the density of the invading immune population can be attributed to the production of immune cells by the dominant population of plasmid bearing cells, CP→CI at a rate y=10^−5^ per cell per hour. If y=0, the density of this immune population would remain at its initial level (“data” not shown). The rate at which the density of immune cells increases is inversely proportional to the fitness cost of carrying the plasmid (Figure S7C). As was the case for where there was no cost associated with the carriage of the plasmid, the increase and leveling off in the density of CI when the plasmid-confers a selective advantage can be attributed to the production of immune cells by the plasmid bearing cells. If not for this, the density of the invading population of immune cells would decline.

**Figure S7.**
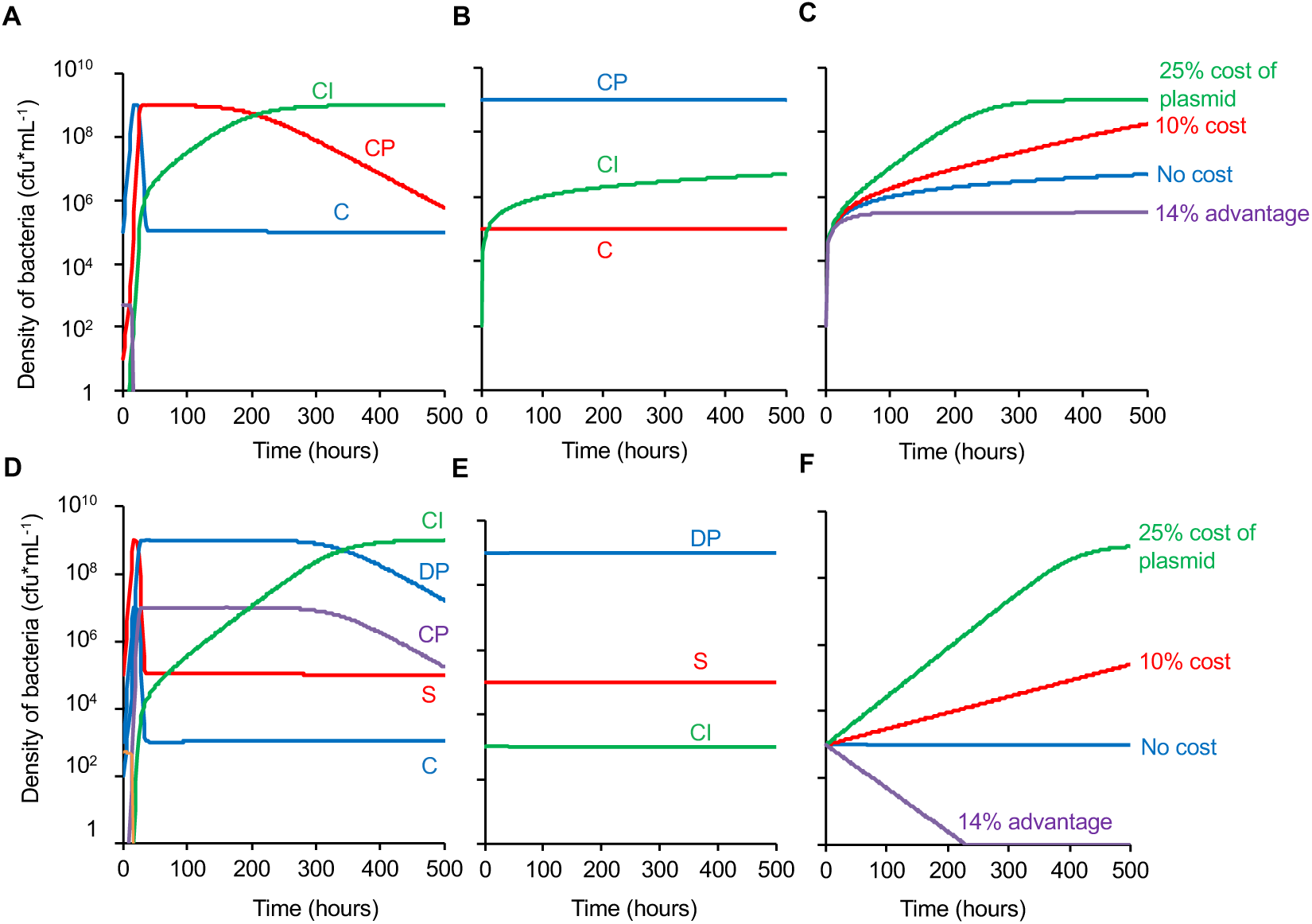
Conditions for the invasion of CRISPR-Cas immunity in populations with conjugative plasmids. Standard parameters, RR=500, w=0.10, e=5×10^−7^, k=1.0, γ=10^−9^, τ=10^−4^, x=10^−4^, y=10^−4^. Changes in the densities of bacteria. A, B and C) Invasion CRISPR-Cas mediated immunity in a CRISPR^+^ population. A) Dynamics of plasmid transfer and the ascent of a CRISPR-Cas immune, CI, population, when there is a 25% cost of plasmid carriage: vC=vCI=0.7, vCP=0.525. B and C) Invasion of CRISPR-Cas immune cells, CI, in population at equilibrium with plasmid bearing and plasmid-free cells, CP. B) Changes in the densities of immune cells, CI, in the absence of selection for or against the carriage of the plasmids: vC=vCP=vCI=0.7.C) Invasion of a CRISPR-Cas immune bacteria, CI, in populations of plasmid-free CRISPR^+^ bacteria with different costs for the carriage of the plasmid: vC=vCI=0.7, vCP=0.7 (no cost), vCP =0.525 (25% cost), vCP =0.63 (10% cost), vCP=1.0 (14% advantage). D-F) Invasion of a CRISPR^-^ population by bacteria carrying CRISPR-Cas immune systems (CRISPR^+^). D) Dynamics of plasmid transfer and the CRISPR^+^ bacteria, CI and CP, in a CRISPR^-^ population with a 25% cost of plasmid carriage: vS=vD=vC=vCI=0.7, vDP=vCP=0.525. E and F) Invasion of CRISPR^+^ cells in a CRISPR^-^ population at equilibrium with the plasmid, DP and S. E) Changes the densities CRISPR^-^ plasmid-bearing and plasmid free cells, DP and S and a CRISPR-Cas-containing population (CRISPR^+^) with plasmid immunity, CI, in the absence of selection for or against the carriage of the plasmid. vS=vDP=vC=vCP=vCI=0.7. F) Changes in the densities of CRISPR^+^ bacteria with different costs of the carriage of the plasmid with different costs for the carriage of the plasmid: vC=vCI=0.7, vCP=0.7 (no cost), vCP =0.525 (25% cost), vCP =0.63 (10% cost), vCP=1.0 (14% advantage).

In Figure S7D, we consider the invasion of CRISPR^+^ cells into a CRISPR^-^ population with a conjugative plasmid. We follow the changes in the densities of the different populations of plasmid-free and plasmid bearing cells in a community initiated with plasmid-free and plasmid-bearing CRISPR^-^ bacteria, S and DP, and a minority population of CRISPR-Cas immune cells, CI. In this simulation, the plasmid engenders of 25% fitness cost and the immune bacteria, CI ascend.

In Figure S7E and F, we consider the invasions of bacteria with CRISPR-Cas immune system into a population of S and DP at equilibrium in the absence of selection for or against the carriage of the plasmid. Under these conditions, the density of the CRISPR-Cas immune cells, CI remains unchanged. If the plasmid engenders a fitness cost, the immune cells invade at a rate that inversely proportional to the cost of the plasmid (Figure S7F).

#### The utility and limits of the models

Richard Levins argued that in constructing mathematical models in population biology one has to, sacrifice generality to realism and precisions, sacrifice realism to generality and precision, or sacrifice precision to realism and generality (147). The models developed here are in this last, heuristic, tradition. Their role is to identify the factors that govern the conditions under which CRISPR-Cas immunity will evolve and the relative contributions of these parameters to this evolution. The parameters of these models can be independently estimated in experimental populations of bacteria and archaea with lytic and temperate phage or conjugative plasmids and the hypotheses generated from their analysis tested in experimental populations of bacteria and archaea with phage and plasmids.

---

**Table S1.** Parameters and Variables.

| **Common Variables and Parameters** | **Symbol and dimensions** | **Value -Range** |
| --- | --- | --- |
| Resource concentration in the habitat | $r \mu\text{g/ml}$ | $0 - 500 \mu\text{g/ml}$ |
| Resource concentration in the reservoir | $RR \mu\text{g/ml}$ | $500 \mu\text{g/ml}$ |
| Volume of the habitat | $Vol$ | $100\text{ml}$ |
| Flow rate into and out of the habitat | $w \text{ ml per hour}$ | $0.10$ |
| $r$ when the growth is half its maximum | $k \mu\text{g/ml}$ | $1.0 \text{ ml}$ |
| Conversion efficiency (resource/cell) | $e - \mu\text{g/cell}$ | $5 \times 10^{-7}$ |
| **Lytic Phage Model** |  |  |
| Density of the phage | $V \text{ particles per ml}$ | $0 - 10^9$ |
| Density of phage sensitive CRISPR<sup>-</sup> | $S \text{ cells per ml}$ | $0 - 10^9$ |
| Density of phage resistant CRISPR<sup>-</sup> bacteria | $SR \text{ cells per ml}$ | $0 - 10^9$ |
| Density of phage sensitive CRISPR<sup>+</sup> bacteria | $C \text{ cells per ml}$ | $0 - 10^9$ |
| Density of phage resistant CRISPR<sup>+</sup> bacteria | $CR \text{ cells per ml}$ | $0 - 10^9$ |
| Density of phage immune CRISPR<sup>+</sup> bacteria | $CI \text{ cells per ml}$ | $0 - 10^9$ |
| Maximum growth rates | $V_S, V_R, V_C, V_{CR}, V_I \text{ hr}^{-1}$ | $0.7$ |
| Adsorption rate constant | $\delta - \text{cells} \times \text{ml/hour}$ | $10^{-7}$ |
| Burst size | $\beta \text{ particles per cell}$ | $50$ |
| Probability of the acquisition of a spacer | $x \text{ per infected cell}$ | $1.67 \times 10^{-8}, 1.67 \times 10^{-7}$ |
| Mutation rates to resistance and sensitive | $\mu_{SR}, \mu_{RS} \text{ per cell per hr}$ | $10^{-8}$ |
| **Temperate phage Model** |  |  |
| Density of free temperate phage | $P \text{ particle per ml}$ | $0 - 10^9$ |
| Density of phage sensitive CRISPR<sup>-</sup> bacteria | $S \text{ cells per ml}$ | $0 - 10^9$ |
| Density of CRISPR<sup>-</sup> lysogenic CRISPR<sup>-</sup> bacteria | $L \text{ cells per m;}$ | $0 - 10^9$ |
| Density of phage sensitive CRISPR<sup>+</sup> bacteria | $C \text{ cells per ml}$ | $0 - 10^9$ |
| Density of CRISPR<sup>+</sup> lysogenic bacteria | $CL \text{ cells per ml}$ | $0 - 10^9$ |
| Density of CRISPR<sup>-</sup> immune bacteria | $CI \text{ cells per ml}$ | $0 - 10^9$ |
| Maximum growth rate | $V_S, V_L, V_C, V_{CL}, V_{CI} \text{ hr}^{-1}$ | $0.525 - 1.0$ |
| Adsorption rate constant | $\delta - \text{cells} \times \text{ml/hour}$ | $10^{-8}$ |
| Burst size | $\beta \text{ particles per cell}$ | $50$ |
| Probability of the acquisition of a spacer | $x \text{ per infected cell}$ | $10^{-5}$ |
| Rate of acquisition of spacers $CP \rightarrow CI$ | $y \text{ per cell per hour}$ | $10^{-5}$ |
| Probability of lysogeny | $\lambda \text{ per infection}$ | $10^{-3}$ |
| Induction rate | $i \text{ per cell per hour}$ | $10^{-4}$ |
| **Conjugative plasmid Model** |  |  |
| Density of CRISPR<sup>-</sup> plasmid free bacteria | $S$ | $0 - 10^9$ |
| Density of CRISPR<sup>-</sup> plasmid-bearing bacteria | $DP$ | $0 - 10^9$ |
| Density of CRISPR<sup>+</sup> plasmid-free bacteria | $C$ | $0 - 10^9$ |
| Density of CRISPR<sup>+</sup> plasmid-bearing bacteria | $CP$ | $0 - 10^9$ |
| Density of CRISPR<sup>+</sup> plasmid immune bacteria | $CI$ | $0 - 10^9$ |
| Maximum growth rates | $V_S, V_{DP}, V_C, V_{CP}, V_{CI} \text{ hr}^{-1}$ | $0.525 - 1.0$ |
| Rate constant of plasmid transfer | $\gamma \text{ cells} \times \text{ml/hour}$ | $10^{-8}$ |
| Segregation rate CP-C, DP->S | $\tau$ per cell per hour | $10^{-4}$ |

